# Human performance in the Traveling Salesman Problem is influenced by spatial scale

**DOI:** 10.1101/2025.10.27.684641

**Authors:** C. Doussot, S. Muflih, J. Mailly, M.M. Müller, N. Boeddeker, M. Lihoreau

**Affiliations:** Univ. Toulouse, CNRS, Research Center on Animal Cognition (CRCA-CBI), Toulouse, France; Bielefeld University, Department for Neurobiology, Germany

**Keywords:** navigation, multi-destination routes, route optimization, Traveling Salesperson Problem, virtual reality

## Abstract

Like many other animals, we humans frequently face complex route optimization problems when planning journeys between multiple locations. Several strategies can be employed to approach such Traveling Salesman Problems. However, these may be strongly constrained by spatial scales, for instance if the goal is to navigate in a supermarket, in a city, or across a continent. Here, we monitored the route optimization performances of human subjects collecting objects in a video game simulating 3D environments at various spatial scales. Unexpectedly, route optimization performances peaked at intermediate spatial scales, where participants could use global optimization strategies. At very large and very small spatial scales, however, participants predominantly employed less efficient local optimization strategies. At the very large scale, the considerable distances between objects prevented them from being seen all at once, making global planning difficult. At the very small scale, participants reported having chosen the shortest path, although they did not. This mismatch between perceived and actual performances suggests suboptimal alternatives were sufficiently short to be considered equivalent to the optimal route. Spatial scale thus strongly influences route planning in human navigators and may determine the spatial behaviors of a wide range of animals facing similar routing problems in their everyday lives.

## Introduction

Animals exploiting patchily distributed resources commonly need to revisit key locations in their environments while minimizing travel costs. Finding the shortest route passing through multiple places is analogous to a formal computational challenge known as the Traveling Salesman Problem (TSP), a NP-hard problem that rapidly becomes intractable(1) as the number of locations increases(1), but for which heuristic algorithms can approach solutions at low computational costs(2).

Humans and many other animals have been described to apply some of these heuristics when asked to solve TSPs by drawing a single stroke linking all dots arranged on a 2D representation (figural TSP)(3–5) or visiting locations across an experimental environment (navigational TSP)(6–8). Local strategies determine the route to follow during only a few steps ahead, like moving to a nearest neighbor target (NN strategy(9)) or grouping visits to nearby targets (Cluster strategy(10, 11)). By contrast, global strategies consider the overall layout and planning of the full route to be used, which generally improves the optimization performance. For example, humans tend to use a convex hull (CH) strategy when presented with figural TSPs(3, 12–14),. They first draw a mental border connecting the outermost targets and gradually pull it inward to include the dots within the polygon shape. This behavior was also observed in monkeys(15), rats(16) and bees(17) in navigational TSPs. However, navigators often appear to combine local and global strategies(5, 18–20) by which first, a coarse approximation of the route is established globally and then refined locally, potentially reducing the cognitive load for the animal(11, 21).

Since TSPs can occur at any spatial scale, distance perception by animals may have a strong impact on the strategies employed and the resulting optimization performances(22, 23). In large-scale environments where target locations are far from each other and potentially hidden by clutter, individuals may rely more on local strategies(7, 24). By contrast, in small-scale environments, where all target locations are visible at once, individuals may use global planning. While the comparison of human performances in a figural TSP (on a piece of paper) and a navigational TSP (in a room-sized environment) suggests individuals use similar heuristics across spatial scales (24), the two tasks involve very different behaviors. They may thus rely on different cognitive mechanisms, including complex visual and attentional processes, as well as spatial memory(22).

Here, we tested the effect of spatial scale on navigational TSP by monitoring the route optimization performances of human subjects collecting 10 objects (chests) in a 3D virtual environment (a Viking village) (25, 26). The participants were asked to repeat the task in the same array of objects but at different spatial scales, so that the distances between the collectibles either decreased or increased while keeping the rest of the visual setting stable. In these conditions, we expected better optimization performances in small environments that allowed global planning, compared to larger environments where local strategies were more practical. Contrary to our expectations, we found that the optimization performance peaked at intermediate spatial scales.

## Results

### Variation in route preferences at three different scales

In a first experiment (Exp1), we asked the participants to collect 10 chests and return to the start location, while taking the fewest steps as possible (**Fig. 1A**). Each participant repeated the task in four arrays of chests: a circular arrangement (one side-bias control), and three experimental configurations (Exp1-C1, C2, C3), at three spatial scales each (S: small (1:2), M: medium, L: large (3:2)) (**Fig. 1D**). In these environments, we measured the utilization of local versus global route optimization strategies by analyzing the first direction (left-right) chosen by the subjects upon leaving the start location N1 (**Table 1**). As expected, we found no side preference by the participants in the control array (Binomial test: corrected p values >0.05 for scale S, M, and L; statistics = 0.60, 0.54, 0.63). However, this was not the case in the three experimental configurations.

**Table 1:**
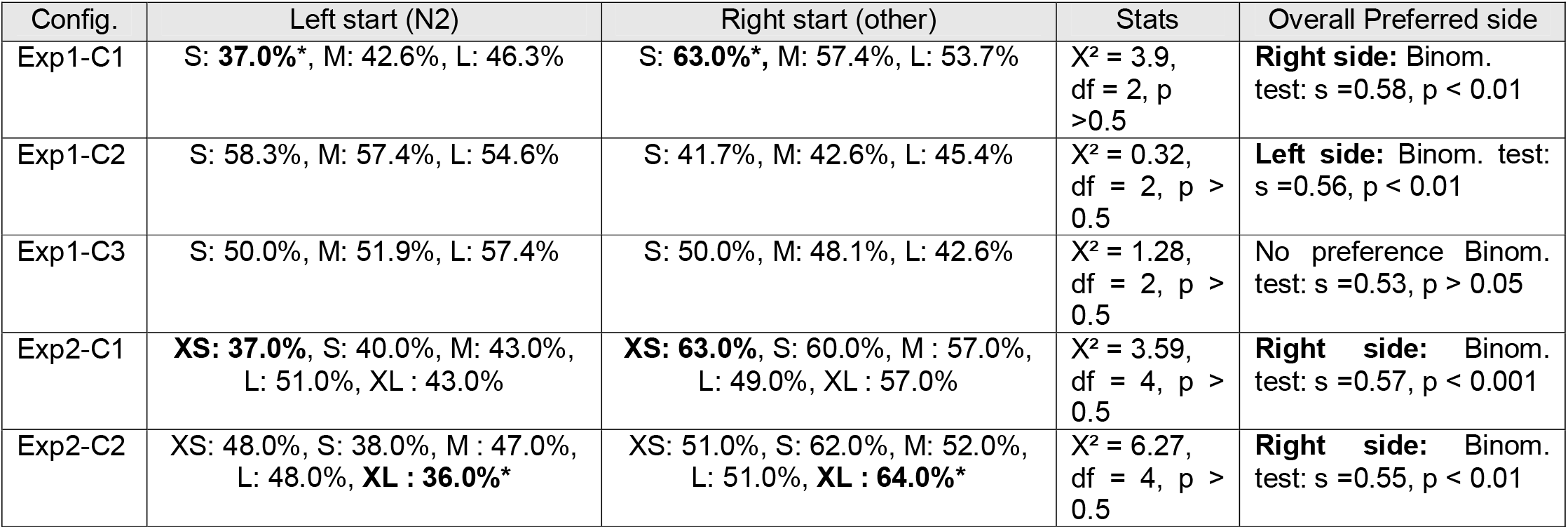
Left - Right preferences across environments and spatial scales. For all conditions, chest N2 is the nearest neighbor from the start location N1. The last column presents Chi-test results assessing side-bias across environments and spatial scales. Significant differences from the binomial test testing for a side preference for each scale are represented in bold. We used 0.05 as the level of significance after Holm-Bonferroni correction at the configuration level. Levels of significance are ***<0.001, **<0.1, *<0.5,. <0.1.

| Config. | Left start (N2) | Right start (other) | Stats | Overall Preferred side |
| --- | --- | --- | --- | --- |
| Exp1-C1 | S: **37.0%\***, M: 42.6%, L: 46.3% | S: **63.0%\***, M: 57.4%, L: 53.7% | $X^2 = 3.9$,<br>df = 2, $p > 0.5$ | **Right side:** Binom. test: $s = 0.58$, $p < 0.01$ |
| Exp1-C2 | S: 58.3%, M: 57.4%, L: 54.6% | S: 41.7%, M: 42.6%, L: 45.4% | $X^2 = 0.32$,<br>df = 2, $p > 0.5$ | **Left side:** Binom. test: $s = 0.56$, $p < 0.01$ |
| Exp1-C3 | S: 50.0%, M: 51.9%, L: 57.4% | S: 50.0%, M: 48.1%, L: 42.6% | $X^2 = 1.28$,<br>df = 2, $p > 0.5$ | No preference Binom. test: $s = 0.53$, $p > 0.05$ |
| Exp2-C1 | **XS: 37.0%**, S: 40.0%, M: 43.0%,<br>L: 51.0%, **XL : 43.0%** | **XS: 63.0%**, S: 60.0%, M : 57.0%,<br>L: 49.0%, **XL : 57.0%** | $X^2 = 3.59$,<br>df = 4, $p > 0.5$ | **Right side:** Binom. test: $s = 0.57$, $p < 0.001$ |
| Exp2-C2 | XS: 48.0%, S: 38.0%, M : 47.0%,<br>L: 48.0%, **XL : 36.0%\*** | XS: 51.0%, S: 62.0%, M: 52.0%,<br>L: 51.0%, **XL : 64.0%\*** | $X^2 = 6.27$,<br>df = 4, $p > 0.5$ | **Right side:** Binom. test: $s = 0.55$, $p < 0.01$ |

**Figure 1:**
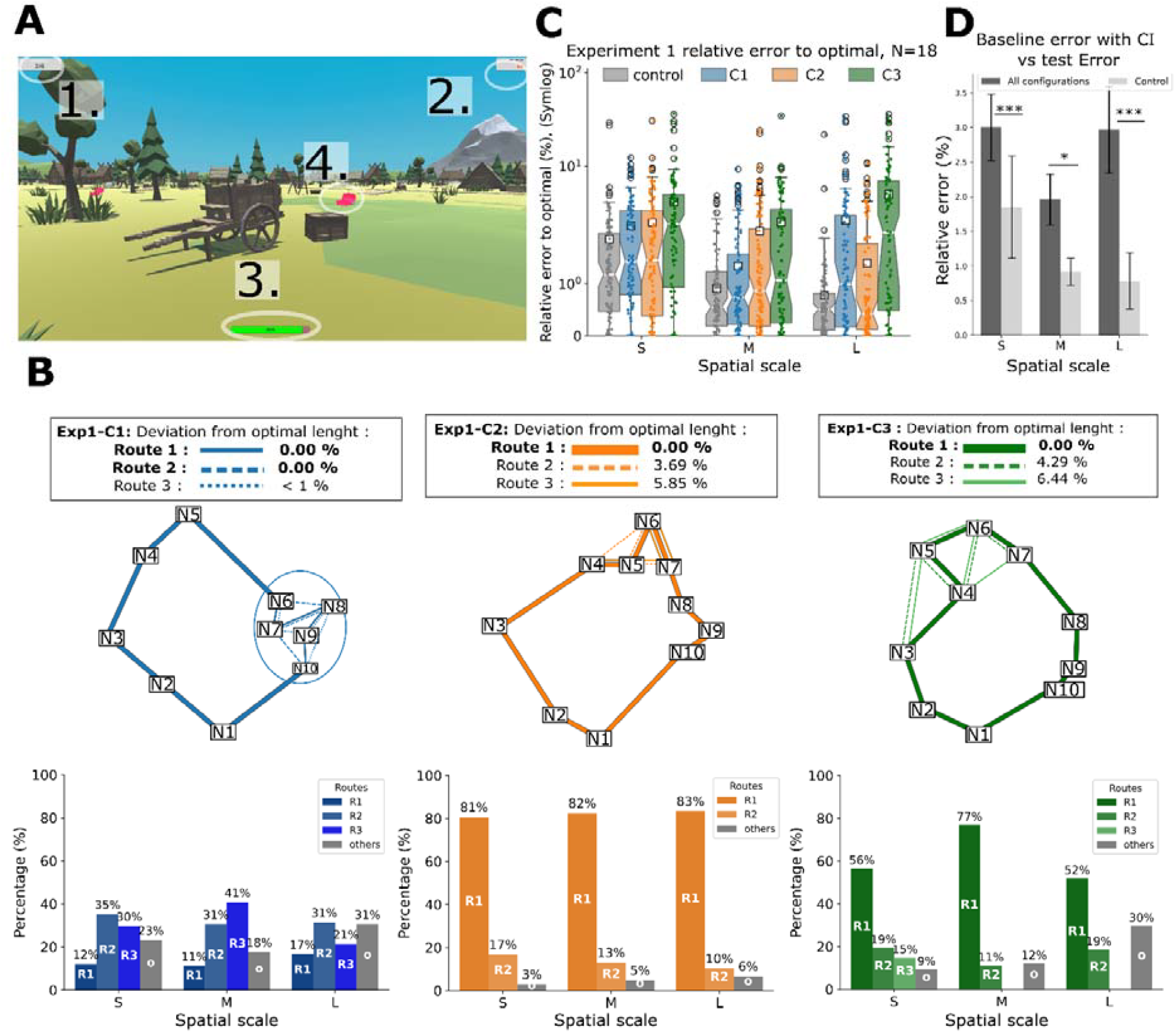
Experiment 1 (N = 18). **A) In-game view**. 1. The inventory indicates the number of chests collected. 2. The number of steps executed from the start. 3. The stamina bar, which decreases as the number of steps increases. 4. Item to collect with a glowing pink texture, indicating it has not been visited yet. **B) Relative error to the optimal trajectory at different spatial scales**. Boxplots of the relative error scaled in symmetric Log due to highly skewed data towards zero. A color represents one configuration (see legend caption). The white squares are the means. **C) Boxplot of Baseline-error distribution (derived from control) versus test-error (C1, C2, and C3) at different spatial scales**. Error bars indicate the Confidence interval (95%). Level of significance between error distributions is indicated with * <0.05, ** <0.01, ***< 0.001. **D) Schematic representation of Configurations 1, 2, and 3** (in blue, orange, and green, respectively): N1-N10 are nodes representing chests. The optimal (R1) and suboptimal topological routes are represented with their associated relative error to optimal %. The color gradient indicates the relative error to optimal, except for Exp1-C1, where all routes are below <1%. Optimal route R1 is the thick solid line, the second-best route the dashed line (R2), and finally the third route the thin line (R3). If more than 4 nodes are close to each other, we regroup them within a circle (cluster). **D) Route choices made by participants in experiment 1**. Proportions of the different routes (sequences of chest collection) observed at the different spatial scales (S, M, L) for the three configurations Exp1-C1, Exp1-C2, and Exp1-C3 from left to right. The colors refer to the routes as defined in the upper graph. Routes performed less than 10% of the time are grouped under the Light Grey bar (and referred to as others).

In the first configuration (Exp1-C1), participants tended to move first to their right (toward chest N10) and collect the cluster of 5 chests (N6-10) rather than moving to the nearest neighbor chest (N2). This preference was the most frequent at the small spatial scale (S) and decreased with increasing scale sizes (**Table 1**). In this configuration, several possible routes (sequences of chest collection) were close to optimal (of less than 1% length difference with the shortest possible route). Participants used a range of these near-optimal routes (R1, R2, and R3). Their frequency of usage varied with spatial scale, suggesting different strategies were used to collect the chests within the cluster (**Fig. 1B**).

In the second configuration (Exp1-C2), participants showed no clear preference between moving left toward the nearest neighbor chest (N2) or right toward a more distant chest (N10; **Table 1**). The optimal route (R1) was preferentially used at all spatial scales (**Fig. 1B**).

In the third configuration (Exp1-C3), participants showed no clear preference between moving left toward the nearest neighbor chest (N2) or right toward two more distant chests (N10 and N9; **Table 1**). This configuration required participants to decide when to integrate the internal chest N4 into their routes. The optimal route (R1) was preferentially use at all spatial scales, and especially at the medium spatial scale (M), where it was observed in 77% of the trials (**Fig. 1B**).

The overall performance of participants, computed from the deviation of their walking trajectories from the shortest possible path (i.e., the relative error) (**Fig. 1D**), shows a complex relationship with spatial scale. Contrary to our expectations, there was no systematic decline in optimization performance with increasing spatial scale. In fact, the opposite was observed in the Control condition as well as in Exp1-C2, where the relative error decreased with increasing spatial scale (GLMM with M scale as reference; β=-0.474, SE=-0.11, t=-4.286, p < 0.001). Responses to spatial scale in Exp1-C1 and Exp1-C3 differed. In Exp1-C1 the relative error shows a quadratic relationship with spatial scale (quadratic component; β=0.486, SE=0.27, t=1.792, p < 0.1), indicating that participants deviated less from the optimal path length at the medium spatial scale. Relative errors in simpler configurations that could be solved by moving in a circle (Control and Exp1-C2), may indicate a baseline level of error attributable to motor imprecision within the video game or an incautious player behavior at a smaller scale. This effect was further magnified because of the normalization process to percentages that we applied. For each spatial scale, the relative error was different from the error-baseline defined by the Control (Mann– Whitney U test; S and L: p < 0.001, M : p < 0.05, and differences in GLMM model estimates see **Table S1**), suggesting that motor imprecision of the participants alone cannot account for the elevated relative error observed, particularly at the smallest and largest spatial scales S and L (**Fig. 1C**). Relative errors also decreased over time as participants gained experience with the game. Across session order, the model revealed a significant reduction in relative error (β = –0.036, SE = 0.013, t = –2.68, p < 0.01) and a significant decrease across trial numbers (β = –0.11, SE = 0.026, t = – 4.13, p < 0.001).

The fact that human subjects performed better at the medium spatial scale by employing more often the optimal route, suggests they changed strategies across scales. However, the three configurations used in our game did not enable us to precisely characterize this change.

### Conflict between global and local heuristics at five different scales

To further explore how human navigators may change optimization strategies across spatial scales, we conducted a second experiment (Exp2) using two new configurations (C1 and C2), in which a direct conflict arose between a global strategy (the convex hull strategy, CH) and one of two local strategies (cluster or nearest neighbor, NN). We enhanced the assessment of performance variation across spatial scales by adding two more scales using the previous M scale as a benchmark (XS, S, M, L, and XL, **Fig. 2A & B**). Moreover, the possible routes inside each configuration were more different from the optimal route than in Experiment 1 (**Fig. 2C**).

**Figure 2:**
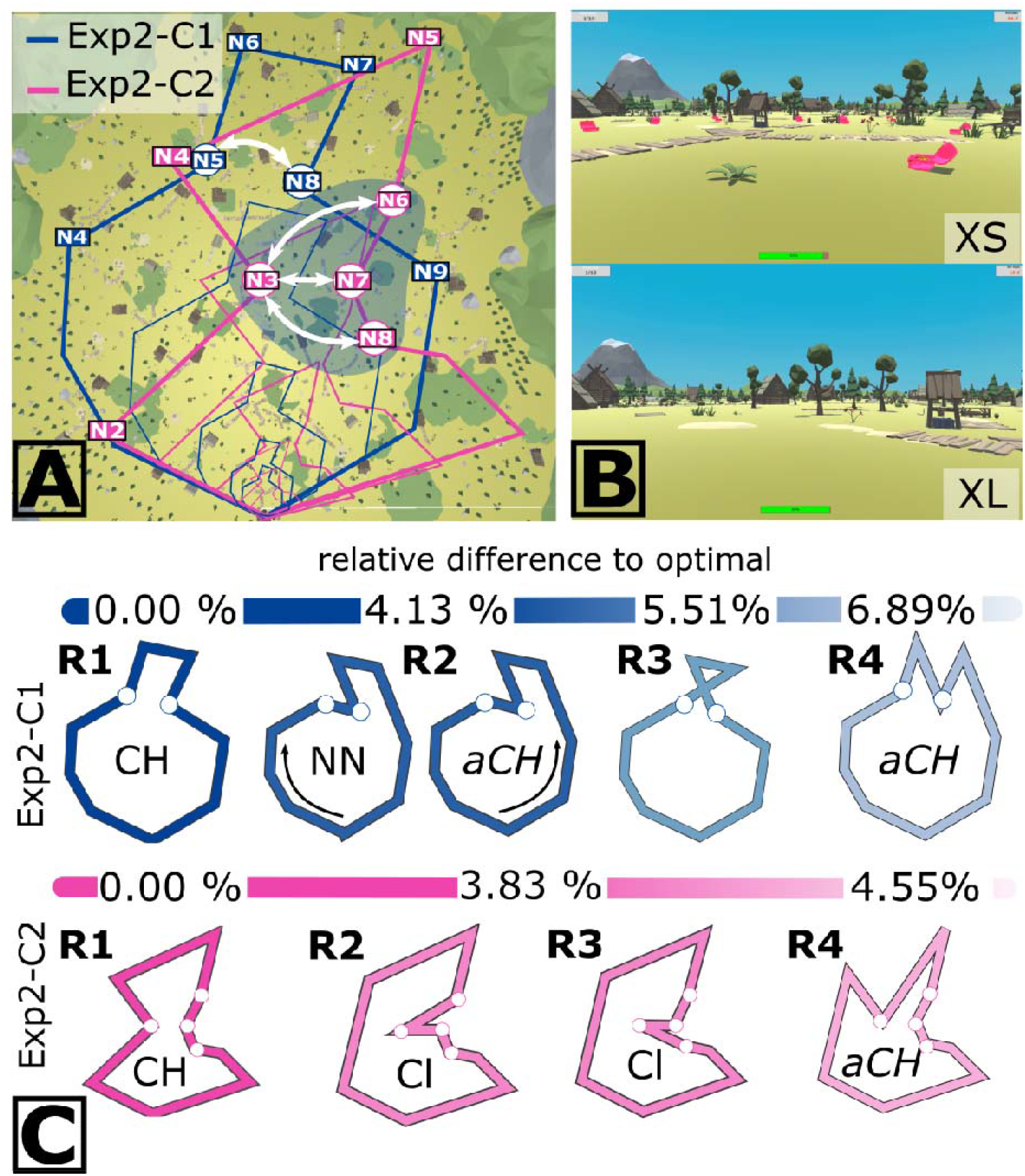
Design of Experiment 2. **A) Game map with all optimal routes** for the two configurations of nodes (chests) at five spatial scales. Broader lines indicate the XL scale. White arrows indicate heuristic conflict locations. For configuration 1 (blue line), there is a possibility for an NN transition between N5 and N8. For configuration 2 (pink line), there is a possibility for a cluster strategy between N3 and any of N6, N7, or N8 (cluster in shaded area). **B) In-game views at the two extreme spatial scales**: Top, smaller scale (XS); bottom, larger scale (XL). **C) Four best routes** for Exp2-C1 and Exp2-C2 and associated optimization strategy. A color gradient shows the percentage difference from the optimal route (R1). Dark blue (optimal) to light blue (non-optimal) for C1, and Pink to light pink for C2. CH stands for convex hull strategy, “aCH” for alternative CH, NN for nearest neighbor strategy, and finally Cl for Cluster strategy.

We designed Exp2-C1 to test the conflict between the usage of an NN and a CH strategy. Participants used the NN transitions between N5–N8 (b) and N7–N9 (e) differently depending on spatial scale. At the small spatial scale S, transitions (b) and (e) were less often used than at the other four spatial scales (N5-N8: X^2^ = 5.83, df = 1, p < 0.05; N7-N9: X^2^= 8.53, df = 1, p < 0.003) (**Fig. 3A**). Corroboratively, the CH transitions showed the opposite trend (N7-N8: X^2^ = 3.32, df = 1, p < 0.1; N8-N9: X^2^= 2.19, df = 1, p > 0.05). To further clarify the strategies employed during transition (b), we differentiated between the two directions: movement from N5 to N8 and from N8 to N5, as the latter does not necessarily reflect an NN strategy but results from initially choosing the most distant chest (N7). Movements from N5 to N8 (i.e., NN strategy) were less frequently observed at medium spatial scales (frequencies: XS: 22.22%, S: 5.75%, M: 8.06%, L:17.24%, ^X^L:12.12%), whereas movements from N8 to N5 (i.e. start with most distant chest) were less frequent at small spatial scales (XS: 6.06%; S: 8.05%; M: 16.13%; L:16.09 %; XL: 14.14%). At spatial scale S, the optimal route (CH strategy) was used most frequently, in 74.4% of the trials (**Fig. 3A**), with R2 as the second most used route (**Fig. 2C**). At the largest spatial scales (L and XL), participants used a more diverse set of routes, with 41–46% of trials involving sub-optimal routes.

**Figure 3:**
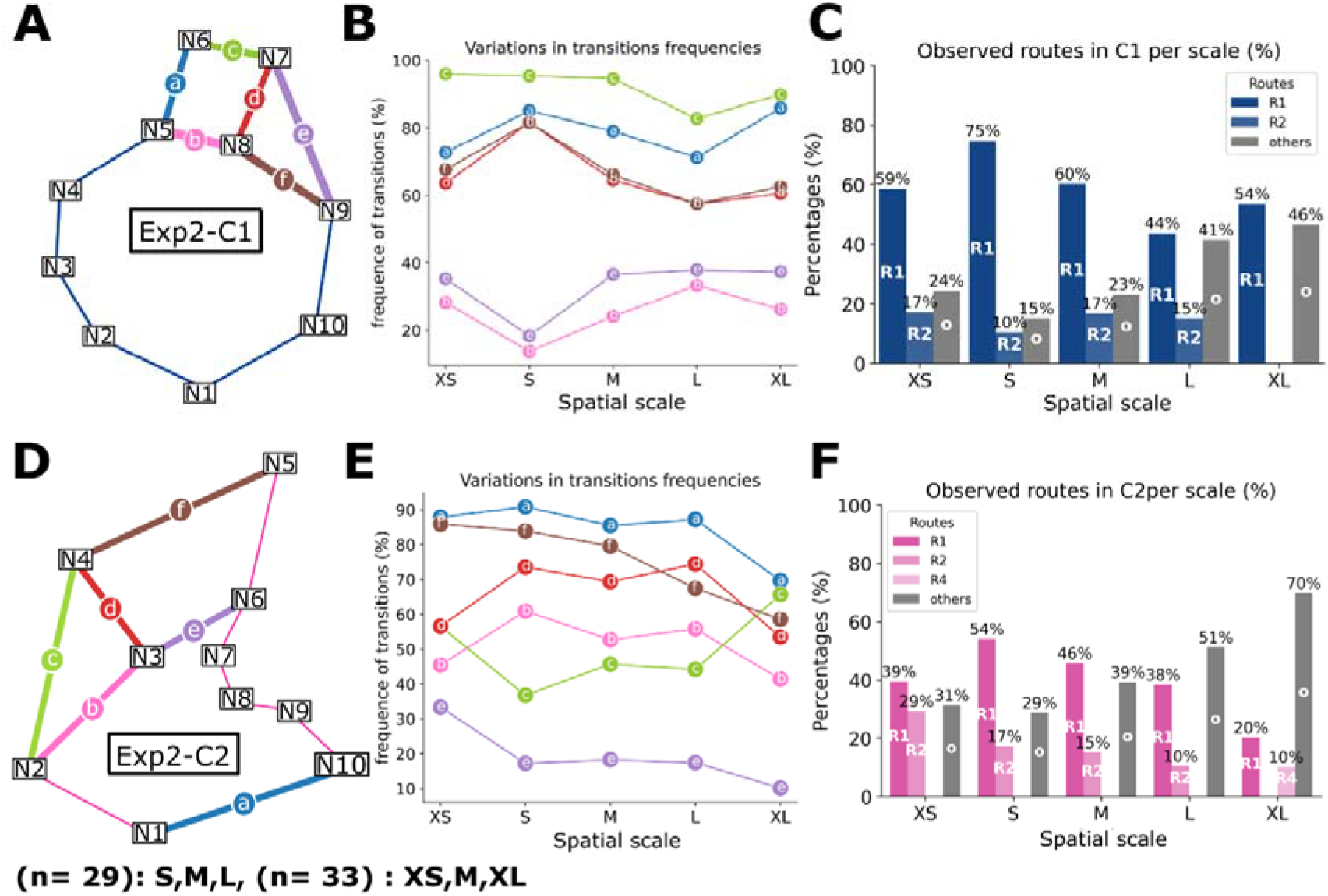
Route choices in Experiment 2. **A) route choice in Exp2-C1**. A is a schematic of the transitions with the highest variability across spatial scales. **B) Occurrence of transitions at each spatial scale**. Letters and associated colors refer to the color scheme in **A. C** is a bar chart of the different routes observed at different spatial scales. Route nomenclature fits the one in Figure 1, and the color scheme the relative error to optimal (from darker optimal to lighter less optimal). **D) route choice in Exp2-C2**. All the subplots **E** and **F** follow the same characteristics of the first row.

In Exp2 - C2, the transition frequencies that varied the most between spatial scales were those created to impose conflict between the CH strategy (R1) and the Cluster strategy (**Fig. 3B**). The transition N2-N4 (c), associated with a misuse of the CH strategy and resulting in clustering the chest N3 with the N6 to N8 chests, was more often used at the two extreme spatial scales (X^2^ = 12.86, df =1, p < 0.001). Occurrence of transitions (b) and (d) varied in consequence of transitions (c). The use of N1-N10 (a), N4-N5 (f), N3-N6 (e) decreased with increasing spatial scale. The optimal route was the most frequently used at medium spatial scales S (54.0% of trials) and M (45.7% of trials) (**Fig. 3B**). Participants used a broader range of routes at large spatial scales: 70% of them were not from the three best ones at the XL spatial scale (**Fig. 3B**).

Across both environments, participants showed a bias to start routes by turning right. In Exp2-C1, this meant they did not preferentially begin by collecting the nearest chest to the start location (N2), while in Exp2-C2, they typically started with the chest that led to the cluster (**Table 2**), which is consistent with the patterns observed in Experiment 1.

**Table 2:**
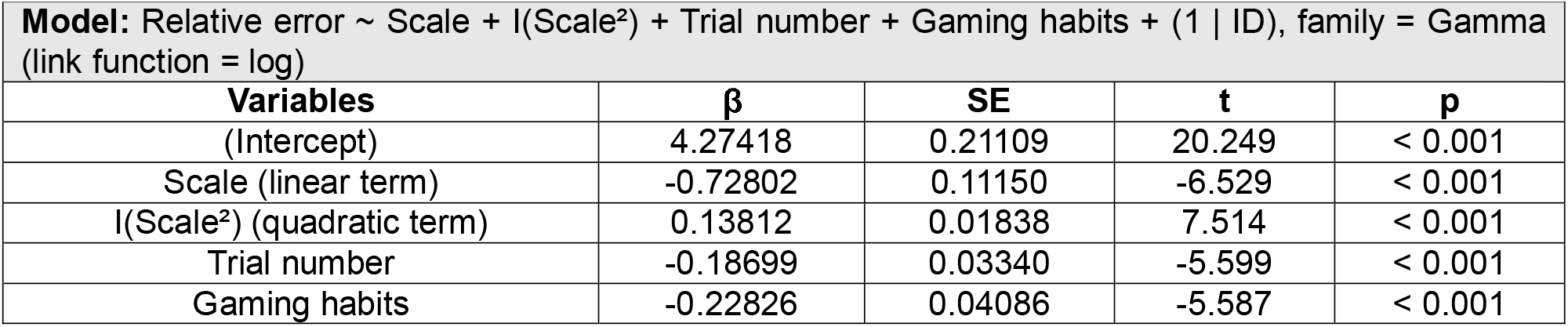
Model predicting the relationship between the relative error from optimal and spatial scale. Model : AIC = 7587.6, BIC = 7622.4. The model follows a quadratic curve, with its linear term representing the scale and the quadratic term being I(Scale^2^), where participants’ IDs serve as a random effect. All predictors are reported in the Variables column. β: estimate, SE: standard error, t: the t-value, and p: p-value.

We then examined whether the scale-dependent pattern observed in the relative error of experiment 1 (**Fig. 1B**) extends to the more extreme spatial scales in experiment 2. The change in spatial scale showed significant predictive power of the error relative to optimal, following a linear component and a quadratic component (**Table 2**). Like in experiment 1, errors peaked at the extreme scales, indicating a quadratic relationship. There was no significant effect of the spatial configuration (C1 and C2). In addition, including inter-individual differences and reducing the number of predictors to gaming habits and trial number in the model increased the marginal R^2^ (see **Table S2** for tested models). Finally, increasing trial numbers and gaming habits significantly reduced the deviation of participants from optimal (**Table 2**; p < 0.001). Our selected model (**Table 2**) was then validated against other models that showed no significance through Likelihood ratio tests and AIC (AIC = 7587.6, X^2^ = 3.7684, df = 3, p = 0.29). Our results and a simplified version of the fitted model are given in **Fig. 4B**.

**Figure 4:**
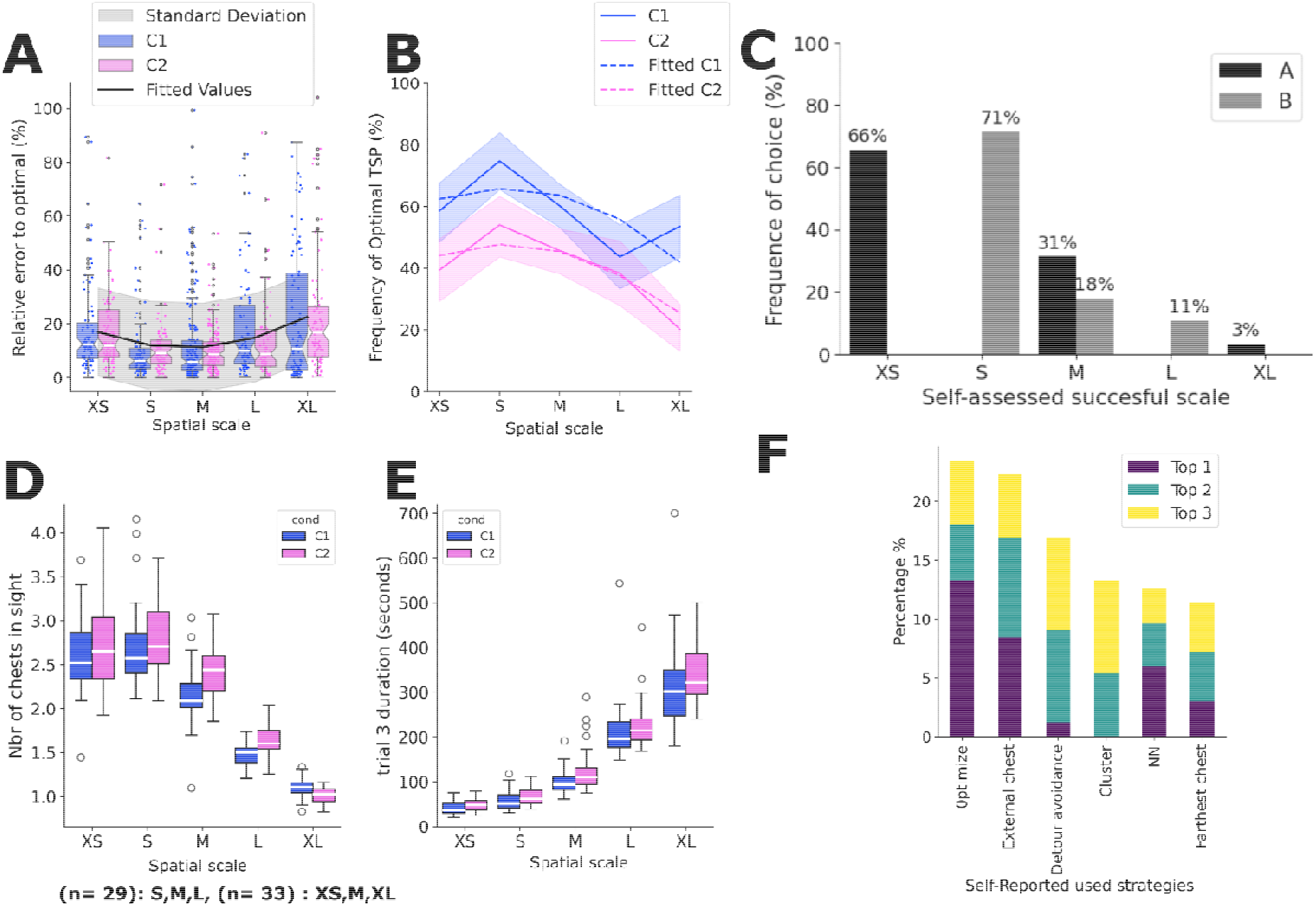
Route optimization performance in Experiment 2. **A) Relative error to the optimal path at different spatial scales**. Colors of boxplots indicate the configurations C1 and C2 (see legend). The median is in white. Dots are individual ata. The black line represents a simplified version of the selected GLMM model (formula = error ∼ scale + I(scale^2)) and the grey shading represents residual deviation. **B) Frequency of optimal route usage** for each configuration with its corresponding standard deviation. The dashed lines represent the fitted values for the simplified version of the selected GLMM (formula = TSP correct ∼ scale + I(scale ^2) + configuration). **C) Proportions of self-assessed use of the optimal route. D)** Average estimated number of chests in sight during the last trial at any scale and configuration. The number of chests in sight was computed based on the participant’s orientation and position in their last trial (trial number 3). **E) Average duration of a game session** in seconds for the different configurations and at each scale at trial number 3. **F) Proportions of self-assessed used strategies**. See in S3 for the detailed formulation of the different criteria present in the feedback survey.

Because Experiment 2 offered clear optimal routes distinct from suboptimal ones, we examined the frequency of optimal route usage by the participants using a GLMM. After the exclusion of unnecessary predicting variables (see **Table S3**), the two chest configurations gave different responses. We compared two models; one in which the relation between the response rate and the spatial scale differed across configurations (with configurations as interaction effect: AIC = 1399.6, BIC 1444.5, df = 9, R^2^ = 18.6%) and one where the performance was lower for Exp2-C2, but the effect of scale remained consistent across configurations (AIC = 1401.3, BIC =1436.2, df = 7, R^2^ = 17.5%). We chose the earlier model because it had fewer degrees of freedom, and the Likelihood-ratio test only suggested a marginal improvement from using the complex model’s prediction (X^2^ = 5.6436, df = 2, p < 0.1).

Spatial scale significantly predicted optimal route usage with a linear (p < 0.1) and a quadratic component (p < 0.01) (**Table 3**). Gaming habits and trial numbers increased the observation of optimal route usage. Participants had significantly higher optimization performance in the first configuration (Exp2 – C1) than in the second one (Exp2 – C2) (p < 0.001, **Table 3**). When accounting for these differences, it increased the conditional R^2^ by 7.7% (Marginal R^2^ = 9.8%, Conditional R^2^ = 17.5%). The distribution of optimal route usage and a simplified version of the fitted model are represented in **Fig. 4B**.

**Table 3:** Model predicting the relation between optimal route usage and the spatial scale. Model: AIC = 1401.3, BIC = 1436.2. The model follows a quadratic curve with its linear term being the scale and the quadratic term being I(Scale^2^) with participants’ ID as a random effect. All predictors are reported in the Variables column. This table reports β: estimate, SE: standard error, t: the t-value and p: p-value

| **Model:** Optimal route observation ~ Scale + $I(\text{Scale}^2)$ + Configuration + gaming habits + trial nb. + (1 \| ID), family = Binomial (logit) | | | | |
| --- | --- | --- | --- | --- |
| Variable | $\beta$ | SE | z | p |
| (Intercept) | -0.65607 | 0.43506 | -1.508 | 0.132 |
| Scale (linear term) | 0.43770 | 0.25566 | 1.712 | 0.087 |
| $I(\text{Scale}^2)$ (quadratic term) | -0.11085 | 0.04203 | -2.638 | 0.008 |
| Conf. C2 | -0.81363 | 0.13225 | -6.152 | < 0.001 |
| Gaming habits | 0.17984 | 0.06332 | 2.840 | 0.005 |
| Trial Number | 0.19449 | 0.08043 | 2.418 | 0.016 |

The results of Experiment 2 thus overall confirmed that the relative error and the optimal route usage follow the same non-linear pattern indicated by Experiment 1 at the extreme spatial scale.

## Discussion

Using a video game, we demonstrated that humans solving a navigational TSP performed better at intermediate spatial scales compared to very small or large ones. This contradicts our initial assumption that global route optimization strategies, associated with higher performances, are primarily used in small-scale environments where all the target locations are visible at once. Instead, we found a complex, scale-dependent relationship between optimization strategies and task performance.

As initially expected, in the video game chest visibility decreased at larger scales (Text S5, LM: β=-0.45, SE = 0.007, t =-61.803, p < 0.001). This hindered global planning and led participants to make incremental decisions based on local considerations, often resulting in longer routes than the optimal one at large spatial scales (**Fig. 4B**). Our result is consistent with the idea that humans are unlikely to represent a sizeable environment within a unique allocentric reference frame(27) and rather use a dynamic representation where remote places are updated intermittently, and nearby ones are continuously maintained(28). Our observations are also in line with reports that the human sense of direction worsens when planning across several regions of interest versus within a single region(29).

However, in contrast to our expectations and despite the good chest visibility, participants also performed poorly on the smallest spatial scales. In these conditions, most participants relied on local strategies, such as favoring the collection of chests placed inside a cluster (Exp2-C2) or moving between nearest-neighbor chests (Exp2-C1). To our knowledge, no study has reported larger errors in distance discrimination and in path integration at a small scale compared to larger ones(30–32), thus calling for alternative explanations. We suggest that at the smallest spatial scales tested in the second experiment, the perception of optimality by the participants becomes ambiguous. In that regard, the surveys filled by the subjects just after being tested are insightful. Variation in perception of optimality can be understood, for instance, in a shopping context when customers do not exclusively reduce their traveled distance but seek to minimize their time carrying heavy weight(33). Similarly, an individual may choose the fastest route over the shortest one: just as a car driver might prefer a slightly longer road with no traffic light to a shorter route with many. In our surveys, participants overestimated their performance for small scales and underestimated it at medium scales (**Fig. 4E**). At the smallest spatial scale (XS), most participants collected chests in a suboptimal sequence and thus exceeded the optimal travel distance. This suggests they did not deliberately settle for “satisficing” (i.e., good enough) solutions(14, 34) or used quicker and less cognitively demanding optimization strategies (i.e., local strategies). Instead, they may have perceived their route as optimal because the task felt quick and feasible, as all chests were visible and routes were overall short. Performance peaked at intermediate spatial scales, reflecting a trade-off: as scale increased, feasibility decreased while the benefit for precise judgment of the optimal route (e.g., saving time) increased. This non-linear pattern mirrors the predictions of resource-rational analysis (35), according to which humans allocate cognitive effort in proportion to the expected gains of optimization (expected utility(36)), leading to maximal performances when cognitive costs and task benefits are balanced. Finally, at the smallest spatial scale, object proximity may have initiated a “crowding-effect”: all items were visible but categorized as equally beneficial(37, 38). Because all distances were short, participants may have considered various routes equally satisfying, despite their actual differences, leading to a choice overload and suboptimal solutions (39). This also raises the question of the extent to which relative distance can be perceived on a smaller scale, a problem that should be specifically addressed in future studies.

Strategy reporting in the surveys further revealed a mismatch between self-perception and actual routing behavior (**Fig. 4F**). If using the optimal route was described as the main strategy by most participants, many reported that they attempted to “*take the external chests to then go to the ones inside*” and “*avoid turns*,”; two behaviors related with the CH strategy(40). Although the NN strategy enabling individuals to follow the optimal path for many transitions, “*going to the closest chest*” was represented in the top three in less than 15% of the surveys. The behavior “c*ollect chests that appear grouped together*”, akin to the cluster strategy, was also underrepresented, given that our results highlight a preference towards node clusters. It is possible that some participants anticipated local strategies to be inherently less efficient for solving TSPs and chose not to report it. This gap between perception and behavior was also noted in an earlier study through analyses of the verbalizations of human subjects tasked to solve the TSP(41). This mismatch could reflect that these strategies may not be “competing” but rather complementary.

Since the discovery of place cells in the rodent brain(42), investigation of navigation’s neural basis often takes place in room-sized environments, where the idea of a perfect mental map(43) can easily be supported. (44–46). Yet, our findings demonstrate that the spatial scale of such tasks is an important factor to consider, as certain scales could be more sensitive to detecting navigation impairments than others. Overall, our data illustrate the importance of accounting for spatial scales when studying spatial cognition. Researchers are now investigating how the neurological components facilitating a cognitive map can be used for larger-scale planning (43). For example, a study suggests a gradual shift from concrete to abstract processing as reflected in brain activity(47), where environments grow beyond the sensory horizon. We argue that navigational TSP is well-suited for studying how neural processes may be at play at different spatial scales during path planning and spatial learning, even outside the human scope(27), like in rodents, birds, and insects. Bees, for instance, are known to optimize their foraging paths when exploiting flower meadows at larger spatial scales than in smaller environments (48, 49). Ultimately such fundamental advances could lead to new applications in the societal domain. For instance, navigation assays based on navigational TSP and fluctuation in spatial scales could help enhance the early detection of neurodegenerative diseases, such as Alzheimer’s disease, where deficits in spatial navigation are among the first cognitive symptoms (44–46).

## Material and methods

### Participants

We ran two experiments with voluntary subjects (see details in **Table 4**). In Experiment 1, we tested 18 healthy participants aged between 19 and 40 years (mean = 29.27 years, SD = 6.17) in 2023. Participants were recruited from the local community through word-of-mouth and acquaintances from Germany and Jordan. In Experiment 2, we tested 62 healthy adults from France, Germany, the UK, and the USA, aged between 18 and 55 years old (mean = 30.27 years, SD = 7.36) in 2024. The participants were selected through word of mouth. We also advertised the project on the university campuses of Toulouse (France) and Bielefeld (Germany). The screening included participants’ demographics: age and self-reported gender identity. Participants were not involved in the study if they reported any untreated mental or visual disorders. All subjects provided informed consent to participate in the game and complete the final survey. They were informed about the nature of the study, the potential risks and benefits, and their right to withdraw from the study at any time. The data collected in the study were anonymized and stored securely. One participant reported that they could not finish the game due to motion sickness; 6 others did not complete the task or made it in several days and were removed to avoid learning biases. The final dataset contained 62 participants. All the participants filled in a survey about personal habits that could influence their performance, including GPS usage, planning practice and video gaming experience(50)

**Table 4:** Methodological details of the two experiments.

|  | Experiment 1 (2023) | Experiment 2 (2024) |
| --- | --- | --- |
| **Number of participants** | 18 | 62 |
| **Duration** | $\cong 3\text{h}$ | 47min11s (SD $\cong 11\text{min}$) |
| **Instructions** | Oral, given by the experimenter | Written on the main menu |
| **stamina bar** | Correlated with scale | Finite: uncorrelated with scale |
| **Number of configurations** | 1 control and 3 tests | 2 tests |
| **Number of spatial scales** | 3 | 5 tested, but only 3 played per participant (game A or B) |
| **Interaction with participants** | Participants were asked to speak out their thoughts while navigating. | No interaction with the experimenter. From home. |

### Game design

For both experiments, we used the same video game created in Unity2021 (Unity Technologies; game available in https://github.com/dousscha/TSP_data_2025). The game interface used to collect the data and create the surveys was made using the Unity Experiment framework toolbox(51), and the game logic was made using the Virtual Navigation Toolbox(52). We used a desktop display similar to previous studies on human navigation(53–56).

The game is in 3D and from a first-person perspective. It requires the use of a mouse and keyboard to control the player’s movements and views. Using a mouse was strongly encouraged; however, two participants used their touchpad in Experiment 2. The game was designed to simulate a navigation task in a village environment (a free Viking village set from the Unity store) that is considered comparable to a “real-world” scenario, in the sense that it offers realistic visual clutter and landmarks (**Fig. 1A**). The main instruction given to players was to collect the treasure chests optimally in the environment while keeping track of the number of steps taken, and to return to the starting location. There were 10 glowing pink chests scattered throughout the environment, but one was positioned directly at the start location. To assess their own performance, the participants were informed that they could check their step count on the top right of the screen, as displayed in **Fig. 1A**. Furthermore, the game interface displayed a stamina bar that ran low with the number of steps taken. In Experiment 2, the virtual stamina decreased at the same rate for all scales to realistically implement the notion of travel cost. When stamina was below 10%, the bar turned red, and a heavy breathing sound was played. This feature was not implemented in the earlier experiment (Experiment 1), where stamina was normalized to the spatial scale investigated (**Table 4**).

We kept the gameplay simple and accessible to a broad audience. The player had to move through the environment with the only constraint that large objects, like houses, could not be crossed. To collect a chest, the player needed to move into its close surroundings (approximately 1 step from its center). The spatial configurations of chests were designed using the center of each chest to compute the optimal route; hence, the circular collection zone around a chest could not be too large, as it would change what route is considered as the TSP solution. When a chest was collected, a sound was displayed, a message appeared on the screen, and the number of chests collected was updated in the left-hand corner of the screen. Furthermore, the color of the chest turned black, but the chest was not removed from the map, so participants would not return to an already visited location. When all the chests were collected, the player was asked through a screen message to return to the start location in order to end the trial. Importantly, the completion of the task was independent of the state of the stamina bar that could be emptied before the player returned to the starting point. During the game, the player’s trajectory, chest collection order, and time to complete the task were saved.

### Study design

#### Experiment 1

We used one control condition (10 chests arranged in a circle) and three spatial configurations (Exp1 C-1, Exp1 C-2, Exp1 C-3) inspired by Blaser and Ginchansky(57) (**Fig. AC**) to test for preferences in optimization strategies at three spatial scales (S, M, and L). The first configuration (Exp1-C1) provided several suboptimal routes whose lengths were close to that of the optimal route (< 1% difference) **(Fig. 1B**). The first and second configurations (Exp1-C1 and -C2) offered a choice between the nearest object (N2) or a cluster of objects (N10) further away from the departure point (N1). In the third configuration (Exp1-C3), participants had to decide when to incorporate the internal node (N4) on their way, following a CH strategy.

These tests were conducted in the presence of an experimenter (see details about instructions in Text S1). Before the test, participants received a brief training session to familiarize themselves with the video game and the TSP navigation task (i.e., a demo session where they had to collect six chests arranged in a circle). The experimenter asked the participant to navigate through the village and collect the objects using the shortest route, using as few steps as possible. Participants were instructed to speak out their thoughts while navigating, to describe the strategies they thought they applied to find the shortest route. The experimenter provided guidance if needed during the experiment. After the demo, participants were randomly assigned a configuration and completed it at all three spatial scales in random order before moving on to the next configuration. They were asked to respond to questions in between each TSP exercise, and another series at the end of the experiment (see questionnaire in Text S2).

#### Experiment 2

Experiment 1 revealed an unexpected quadratic spatial scale effect on the global performance of participants and affected their route choices (**Fig. 1)**. We therefore performed a complementary Experiment 2 with two new configurations (Exp2-C1, Exp2-C2), conflicting global and local strategies, and at more spatial scales (XS, S, M, L, XL) to include extremely small and large ones (**Fig. 2A)**.

Exp2-C1 was used to test the CH and NN strategy. In this configuration most transitions composing the optimal route followed NN transitions, except the transition between N5 and N8, which belonged to the CH route (**Fig. 2A**). At this location, using the shortest transition led to a suboptimal solution of the TSP (R2) (**Fig. 2C**). Yet in R2, moving from N8 to N5 might not be a NN transition if it results from moving to N7 from N9 first; in this case the participant may decide to collect first the farthest node or attempt a CH strategy, referred as an alternative CH (aCH) (**Fig. 2C**). The best suboptimal route (R2) was 4.13% longer than the optimal.

Exp2-C2 had a cluster of 5 chests positioned on the right of the departure location (**Fig. 2A**). According to previously mentioned strategies, the player could either collect all the chests together using the cluster strategy (R2 and R3) or include the outermost chest of the cluster (N3) while forming the left border of the convex shape (CH strategy) (**Fig. 2A**). Performing the latter would lead to the optimal route (R1). The best suboptimal route among all possibilities following the cluster strategy would lead to a 3.83% deviation from the optimal (R2).

Based on the results of experiment 1, where optimal routes were most observed at a medium spatial scale, we tested participants on five spatial scales (326 steps in length). The medium spatial scale was 363 steps long for Exp2-C1 and 418 for Exp2-C2, and the four other spatial scales were two to three times smaller or longer, ranging from XS to XL (**Fig. 2A&B**). To reduce experimental time and the risks that participants abandon the task, they were divided into two groups, each tested on three of the five spatial scales rather than all five. One group of participants was tested at spatial scales XS, M and XL (N = 33) and was asked to download the game A, and the other group played at scales S, M and L (N= 29) in game B. To control for potential learning effects on the TSP performances, the order in which the participants performed the scale was pseudo-randomized. The participants systematically started with Exp2-C1 followed by Exp2-C2 at the same spatial scale. They all had three trials per session (configuration and scale). In total, each of them did 6 sessions, taking on average a total of 47 minutes and 11 seconds (SD,… 12 minutes).

Experiment 2 was performed without the supervision of an experimenter. The participants received an email with a link to download a game, A or B, depending on the group they were assigned. The invitation included all the necessary details to run the game: the ID used for later anonymization, the order in which to play the mini games (i.e., a specific combination of configuration and spatial scale, unknown to the player), and a notice to send their data back. The email informed the participants of approximate duration of the experiment (between 50 and 65 minutes) so that they could plan their time accordingly. Upon launching the game, the menu screen indicated the aim of the game, the consent form, and some of the pre-experiment survey questions (see questionnaire in **Text S3**). These questions were asked beforehand to avoid biasing the results due to the experience of playing the game (see full menu screen text in **Text S4**). The players run the demo first, to familiarize themselves with the setup. After, participants played each mini game in the order they were given. At the end, the participants filled out a final survey about their optimization routine and performance assessment (see questions in S3). The questions followed either a Likert-based or a semantic differential-based scoring system(58). The game was available in French and English.

### Data analysis

The raw data are available in https://github.com/dousscha/TSP_data_2025.

We studied left-right biases in the routes of participants to explore strategy preferences in Experiment 1 and side preference in Experiment 2. We used an exact binomial-test to test for preferences and a chi-squared test to identify changes in proportions of strategy/side usage due to spatial scale. We further characterized the routes taken by participants across scales in both experiments, focusing on the transitions between chests. To assess variation in frequencies of transition between two chests, we applied chi-squared tests.

We then looked at various metrics to describe the participants’ performance. Firstly, for both experiments, we measured the relative length deviation between each participant’s trajectory and the optimal path (i.e. relative error, F1).

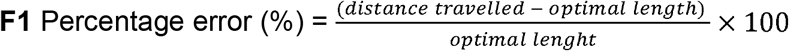

The optimal route, i.e., the shortest possible route to collect the 10 chests, was first computed based on the route passing through each chest center (i.e., theoretical optimal). However, due to the virtual catchment zone around each chest allowing their virtual collection, the actual optimal path length might be slightly shorter than the computed one. Consequently, the optimal was recomputed from the minimum of all candidates if this one happened to be below our theoretical optimal.

Secondly, and only for Experiment 2, because it offered an optimal route that was clearly different from the suboptimal ones (3.83% difference minimum, **Fig. 2**), we reported success in optimally solving the TSP for each trial, i.e., collecting the chest in the optimal order independently of the walked trajectory. Contrary to the error, this metric does not account for any zigzagging movements that could be due to poor gaming habits.

We analyzed the data using Python and R. For the performance metrics (relative error and optimal TSP frequency), we fitted GLMMs in R. Initial models included effects of trial number, session order, age, gender, gaming habits, GPS usage, navigation assessment, and optimization practice. Non-predicting variables were removed during model selection. Player ID was also included as a categorical random effect to examine for inter-individual differences. We always tested for the interaction effect of the configurations and the significant predictive variable. Models were selected based on the Akaike Information Criterion (AIC), Likelihood ratios test, and their simplicity (i.e., low degree of freedom). Hence, interactions were removed since they were shown not to be significant and worsen the model simplicity. When necessary, due to the non-linearity of the variables’ responses (relative error and optimal TSP solving frequency), we used the spatial scale as a numeric predictor during model fitting, even though we did not assume equidistance in the response relative to increasing scale. We used a gamma distribution with a Log link function for the relative error. While the success of optimal TSP in a trial was either 0 (no optimal TSP) or 1 (optimal solving), we used a binomial distribution for model fitting. To allow for proper model fitting and analysis, outliers were removed using the z-score metric; any results with a z-score above three were removed.

## Ethics statement

All experiments were conducted in accordance with the guidelines of the Deutsche Gesellschaft für Psychologie e.V. (DGPs) and approved by the Bielefeld University Ethics Committee. The studies were conducted in accordance with the local legislation and institutional requirements. The participants provided their written informed consent to participate in this study.

## Author contributions

CD: Conceptualization, Data curation, Formal analysis, Methodology, Software, Visualization, Writing—original draft, Writing—review & editing. SM: Conceptualization, Methodology, Data curation, Writing—review & editing, JM: Methodology, Writing—review & editing. MM: Software, Writing—review & editing. NB: Project administration, Supervision, Writing—review & editing. ML: Methodology, Funding acquisition, Project administration, Resources, Supervision, Writing—review & editing.

## Acknowledgments

We thank all participants for their time and Renaud Bastien for early comments on the manuscript.

