## Supplementary Materials for "Human performance in the Traveling Salesman Problem is influenced by spatial scale"

**This document contains:**

Text S1: Supplementary methods for Experiment 1

Text S2: Questionnaire used in Experiment 1

Text S3: Questionnaire used in Experiment 2

Text S4: Main menu screen with game instructions

Text S5: Computation of chest visibility

Table S1: GLMM model fitting on relative error for Experiment 1

Table S2: GLMM model fitting on relative error for Experiment 2

Table S3: GLMM model fitting on TSP optimal success for Experiment 2

**Text S1: Supplementary methods for Experiment 1**

Participants training and Procedure:

The experiment was performed in the presence of a researcher. Prior to the start of a test, participants were given a brief training session to familiarize themselves with the video game and the navigation task (the demo). The participants were instructed to speak out their thoughts while completing the navigation task, describing the strategies they think they apply for finding the shortest possible route in the virtual environment. The experimenter instructed the participants to: “*Navigate through the village and collect the glowing objects (always ten objects, except the demo trial: six objects) using the shortest possible route, to make it clearer; using as few steps as possible, where you can check your performance by checking the steps count (top right). The starting point, where the first object is already automatically collected, is also the finishing point of the trial*”. The researcher provided feedback and guidance to the participants as needed during this familiarization phase.

Once the participants were comfortable with the navigation task, they began the experiments. Each trial began with a reminder of the task instructions and a reminder to verbalize their thoughts. The participants were randomly assigned one of the four configurations, completing three trials within the same configuration, each on a different scale. The order of the scales was randomized as well. After completing all three sessions for a given configuration, the participants were permitted to take a short break since the experiment could run for up to three hours.

**Text S2: Questionnaire used in Experiment 1**

After each trial

Trial: Configuration-spatial scale / trial number:

-	Describe in two sentences the strategy you used to find the shortest route in this exercise.

-	How comfortable was this exercise on a scale of 10.

After the experiment (all trials completed)

- How frequently do you play video games on average?

- Do you usually try to plan your route in your daily activities?

- Do you usually try to optimize your routes in your daily activities?

- How often do you navigate with a GPS?

**Text S3: Questionnaire used in Experiment 2**

Before the experiment

- **Gender:** Male, Female, Other, Do not wish to answer*.*
- **Age/*Age:***
- **You play video games/*Vous jouez aux jeux videos:*** Never, Once a month or less 2 to 4 times per month, 2 to 3 times per week, 4 times or more per week.
- **You are good at orienting/*Vous êtes bon.ne en orientation*:** Completely disagree, Disagree, Rather Disagree/, Rather Agree, Agree, Completely agree.
- **You use a GPS/ *Vous utilisez un GPS*:** Never, Rarely, occasionally, regularly, daily.

After the experiment

- **To play the game, did you use a mouse*?***
- **What was your average impression of the difficulty of the exercise (interface and aim of the game combined)?** Very easy, Easy, Manageable, Hard*,* Very hard*.*
- **On average you think you chose the shortest path linking all chests.** Completely disagree, Disagree, Rather Disagree, Rather Agree, Agree, Completely agree.
- **Do you think you succeded best at finding the optimal route at small or at large spatial scale**Small, Medium*,* Large*,* Cannot answer.
- **Select all the strategies you used when deciding on which chest to collect :** A) Go to the closest chest, B) Take the chest further away and then get the others, C) Take the external chests to then go to the ones inside*,* D) Take the most visible chests, E) Avoid detour (limit turns), F) Collect chests that appear grouped together, G) Optimize the global path*.*
- **From the list above, choose the three strategies you used the most on average, in order from the most used to the least (in this notation: A,B,C)*.***
- **In your everyday life, you optimize your journeys ?** Completely disagree, Disagree, Rather Disagree, Rather Agree, Agree, Completely agree.

**Text S4: Main menu screen with game instructions**

Read the instructions and give your consent:

Press "Esc" to pause at any moment.

Important:

Play all the levels in the order given in the invitation. Begin with the demo, there are three trials.

To launch the game: Manually select in the drop-down menu the next level at the end of each one (1). Select the folder "Results_Game_A (or B)" located in the game’s downloaded folder (A or B) (2). Write down your given ID (3), then after each new level, record the completion order, starting with 1, then 2, and so on until 7 (4), starting with the demo as “1.” Then answer the questions.

Objective of the game: Find all the chests in the village and optimize their collection. Keep an eye on your energy bar at the bottom of the screen, which decreases with each step you take. You can only return to the starting point once all 10 chests have been collected. The first chest is located at your starting point, ensuring that you won't go back empty-handed. Once the chests are gathered, deposit them at the starting point by pressing the E key. You will need to complete this task 3 times per game.

Controls: Use the arrow keys on your keyboard to walk in the environment OR press W to move forward, A to go left, D to go right, and S to move backward. Use the mouse to adjust the camera and look around. Small objects can be walked through, so you won't need to go around them.

**Text S5: Computation of chest visibility**

We used the Unity ray-tracer to assess occlusions from the houses and estimate players’ views by defining a horizontal field of view of $\cong$ 90°(following the most common screen aspect-ratio (16:9) based on their orientation during the last trial of each session. Thus, selecting only their best performance as the trial number is positively correlated with performance. Chests became barely visible when moving further than 173 steps from them (threshold defined manually). With all the information, we postprocessed at each location of their down-sampled trajectories (down-sampling rate: 12, 18, 37, 75, 112 for XS, S, M, L and XL, respectively) the number of chests that are visible at each instant.

**Table S1: GLMM model fitting on Relative error for Experiment 1**

Model : Relative error ~ num_scale * Configuration + I(num_scale^2) *Configuration

+ session nb + trial nb +(1|player ID), family=Gamma(link = "log")

| Term | Estimate (M) | Std. Error | t value | Pr(>\|t\|) | Significance |
| --- | --- | --- | --- | --- | --- |
| (Intercept) | 0.65925 | 0.21106 | 3.124 | 0.00183 | ** |
| num_scale | -0.47381 | 0.11055 | -4.286 | 1.97e-05 | *** |
| ConfigurationC1 | 0.23104 | 0.22178 | 1.042 | 0.29773 |  |
| ConfigurationC2 | 0.67769 | 0.22053 | 3.073 | 0.00217 | ** |
| ConfigurationC3 | 1.07731 | 0.22343 | 4.822 | 1.61e-06 | *** |
| I(num_scale^2) | 0.12879 | 0.19113 | 0.674 | 0.50055 |  |
| trial | -0.11037 | 0.02672 | -4.130 | 3.88e-05 | *** |
| session_num | -0.03619 | 0.01350 | -2.680 | 0.00746 | ** |
| num_scale:ConfigurationC1 | 0.47914 | 0.15674 | 3.057 | 0.00229 | ** |
| num_scale:ConfigurationC2 | 0.13572 | 0.15732 | 0.863 | 0.38846 |  |
| num_scale:ConfigurationC3 | 0.50857 | 0.15953 | 3.188 | 0.00147 | ** |
| ConfigurationC1:I(num_scale^2) | 0.48571 | 0.27099 | 1.792 | 0.07334 | . |
| ConfigurationC2:I(num_scale^2) | -0.12470 | 0.27088 | -0.460 | 0.64536 |  |
| ConfigurationC3:I(num_scale^2) | 0.32468 | 0.27447 | 1.183 | 0.23708 |  |

**Table S2: GLMM model fitting on Relative error for Experiment 2**

*Model 1*: AIC = 7681.8, BIC = 7766.2

*formula = response ~ Scale * Configuration + I(Scale²) * Configuration + Trial + Gaming habits + GPS usage + daily optimization + session + Age + Gender + navigation rating, family = Gamma (link = "log").*

| Variables | Estimate | Std. Error | t value | Pr(>\|t\|) |
| --- | --- | --- | --- | --- |
| (Intercept) | 4.261163 | 0.378472 | 11.259 | < 2e-16 *** |
| Scale | -0.575855 | 0.157037 | -3.667 | 0.000258 *** |
| Conf C2 | 0.329853 | 0.301581 | 1.094 | 0.274318 |
| I(Scale²) | 0.113215 | 0.025580 | 4.426 | 1.06e-05 *** |
| Trial number | -0.157237 | 0.037956 | -4.143 | 3.71e-05 *** |
| Gaming habits | -0.195009 | 0.025377 | -7.684 | 3.53e-14 *** |
| GPS usage | -0.033635 | 0.036634 | -0.918 | 0.358766 |
| Daily optimisation | -0.073494 | 0.031096 | -2.363 | 0.018288 * |
| Session nbr | -0.018648 | 0.018912 | -0.986 | 0.324328 |
| Age | 0.013485 | 0.004846 | 2.783 | 0.005489 ** |
| Gender autre (n=1) | 0.317762 | 0.264877 | 1.200 | 0.230545 |
| Gender female | -0.028827 | 0.077731 | -0.371 | 0.710822 |
| Gender ne souhaite pas répondre (n=1) | -0.733402 | 0.263745 | -2.781 | 0.005521 ** |
| Navigation rating | -0.022955 | 0.030160 | -0.761 | 0.446759 |
| Scale:condC2 | -0.272152 | 0.219219 | -1.241 | 0.214714 |
| Cond C2:I(scale²) | 0.041911 | 0.035704 | 1.174 | 0.240717 |

*Model 2 :* AIC = 7589.8 BIC = 7639.5

*formula = response ~ num_scale + I(num_scale^2) + Trial + Gaming habits + GPS usage + daily optimization + Age + (1 | Participants ID)*

| Variables | Estimate | Std. Error | t value | Pr(>\|z\|) |
| --- | --- | --- | --- | --- |
| (Intercept) | 4.197454 | 0.479245 | 8.758 | < 2e-16 *** |
| num_scale | -0.717228 | 0.111552 | -6.430 | 1.28e-10 *** |
| I(num_scale^2) | 0.136401 | 0.018391 | 7.417 | 1.20e-13 *** |
| trial_num | -0.186683 | 0.033428 | -5.585 | 2.34e-08 *** |
| gaming_habits | -0.202126 | 0.041986 | -4.814 | 1.48e-06 *** |
| GPS_uage | -0.012360 | 0.063871 | -0.194 | 0.847 |
| daily_optimisation | -0.082858 | 0.054084 | -1.532 | 0.126 |
| Age | 0.013389 | 0.008903 | 1.504 | 0.133 |

**Table S3: GLMM model fitting on TSP optimal success for Experiment 2**

*Model 1:* AIC = 1415.6, BIC = 1416.1

formula = TSP_correct ~ num_scale * cond + I(num_scale^2) *

cond + trial_num + gaming_habits + GPS_uage + daily_optimisation +

session_num + Age + Gender + +navigation_rating, family = binomial(link = "logit")

| Variables | Estimate | Std. Error | z value | Pr(>\|z\|) |
| --- | --- | --- | --- | --- |
| (Intercept) | -0.472224 | 0.785349 | -0.601 | 0.54765 |
| num_scale | -0.005274 | 0.322672 | -0.016 | 0.98696 |
| condC2 | -1.717748 | 0.625953 | -2.744 | 0.00607 ** |
| I(num_scale^2) | -0.029586 | 0.052270 | -0.566 | 0.57137 |
| trial_num | 0.186745 | 0.078717 | 2.372 | 0.01768 * |
| gaming_habits | 0.139489 | 0.052188 | 2.673 | 0.00752 ** |
| GPS_uage | 0.097045 | 0.077199 | 1.257 | 0.20873 |
| daily_optimisation | -0.014047 | 0.064924 | -0.216 | 0.82870 |
| session_num | 0.007655 | 0.039341 | 0.195 | 0.84571 |
| Age | -0.001740 | 0.009977 | -0.174 | 0.86154 |
| Genderautre | -0.307096 | 0.560282 | -0.548 | 0.58362 |
| Genderfemale | -0.299326 | 0.158383 | -1.890 | 0.05877 . |
| Genderne souhaite pas répondre | 1.830033 | 0.682418 | 2.682 | 0.00733 ** |
| navigation_rating | 0.073934 | 0.062441 | 1.184 | 0.23639 |
| num_scale:condC2 | 0.887176 | 0.456855 | 1.942 | 0.05215 . |
| condC2:I(num_scale^2) | -0.162089 | 0.075353 | -2.151 | 0.03147 * |

Model 2: AIC = 1399.6, BIC = 1444.5

Formula = TSP_correct ~ num_scale * cond + I(num_scale^2) * cond + gaming_habits +

trial_num + (1 | ppid)

| Variables | Estimate | Std. Error | z value | Pr(>\|z\|) |
| --- | --- | --- | --- | --- |
| (Intercept) | -0.16810 | 0.53983 | -0.311 | 0.75550 |
| num_scale | -0.02091 | 0.34606 | -0.060 | 0.95182 |
| condC2 | -1.82795 | 0.64396 | -2.839 | 0.00453 ** |
| I(num_scale^2) | -0.02871 | 0.05616 | -0.511 | 0.60919 |
| gaming_habits | 0.18080 | 0.06367 | 2.840 | 0.00452 ** |
| trial_num | 0.19547 | 0.08064 | 2.424 | 0.01535 * |
| num_scale:condC2 | 0.95446 | 0.47002 | 2.031 | 0.04229 * |
| condC2:I(num_scale^2) | -0.17380 | 0.07748 | -2.243 | 0.02489 * |
